# Using strain-resolved analysis to identify contamination in metagenomics data

**DOI:** 10.1101/2022.01.16.476537

**Authors:** Yue Clare Lou, Jordan Hoff, Matthew R. Olm, Jacob West-Roberts, Spencer Diamond, Brian A. Firek, Michael J. Morowitz, Jillian F. Banfield

**Author notes:** Correspondence (J.F.B.). Department of Microbiology and Immunology, Stanford University School of Medicine, Stanford, CA 94305, USA.

## Abstract

Metagenomics analyses can be negatively impacted by DNA contamination. While external sources of contamination such as DNA extraction kits have been widely reported and investigated, contamination originating within the study itself remains underreported. Here we applied high-resolution strain-resolved analyses to identify contamination in two large-scale clinical metagenomics datasets. By mapping strain sharing to DNA extraction plates, we identified well-to-well contamination in both negative controls and biological samples in one dataset. Such contamination is more likely to occur among samples that are on the same or adjacent columns or rows of the extraction plate than samples that are far apart. Our strain-resolved workflow also reveals the presence of externally derived contamination, primarily in the other dataset. Overall in both datasets, contamination is more significant in samples with lower biomass. Our work demonstrates that genome-resolved strain tracking, with its essentially genome-wide nucleotide-level resolution, can be used to detect contamination in sequencing-based microbiome studies. Our results underscore the value of strain-specific methods to detect contamination and the critical importance of looking for contamination beyond negative and positive controls.

## Introduction

The advancement of sequencing technologies has enabled researchers to investigate microbial communities at high resolution and throughput. However, contamination poses challenges for data analysis. Contamination refers to DNA within a sample that did not originate from that specific biological sample. Recognizing contamination, followed by appropriate decontamination, should be a critical first step for all microbiome analyses. Skipping this step can easily result in confounded results and false conclusions.

Contamination can originate outside of a study. Microbial DNA from the surrounding environment, native microbiomes of researchers, and microbial DNA present in DNA extraction and library preparation kits are all considered external contaminants [1–3]. Detection of such contamination is enabled by the addition of negative controls (i.e., blank reagent controls) during sample collection, preparation and/or sequencing. To date, strategies have been devised to minimize, detect and/or remove externally derived contamination *in silico* [1,3–6].

Cross-sample contamination is less well studied [3,7]. Contaminants that originate within a study can be introduced during DNA extraction [7] when DNA from one sample spills over into another. It can also occur during sequencing as a result of index switching or sample bleeding [8,9]. Since the contaminating DNA in this case originates from microorganisms present in samples from the study, one cannot decontaminate by removing “contaminant species” present in controls from the actual dataset. While strategies have been developed for solving index switching and sample bleeding arising during sequencing [9,10], much less is known about the well-to-well contamination occurring prior to sequencing in metagenomics data and how to detect it.

In recent work that characterized early-life intestinal strain colonization, we collected a large clinical dataset consisting of over 400 fecal samples from infants and their mothers [11]. Unfortunately, we detected clear evidence of cross-sample contamination that required us to exclude one half of the samples from the study. This motivated an in-depth investigation of the signals used to detect contamination in this and a second clinical dataset of preterm infants. Our approach relied upon a strain-resolved workflow. Strain-level analyses have been increasingly applied in microbiome studies, for example, to study mother-infant gut microbiome transmission [11–13] and conduct epidemiological surveillance [14,15]. The high specificity and sensitivity of strain tracking methods is powerful, yet it is also clear that conclusions from these methods can be confounded by cross-sample contamination. Here, we provide a framework for the detection of cross-sample contamination using strain-based surveillance, and in two case studies, we show how such a framework can be used to detect contamination in large-scale metagenomics datasets.

## Results

### Case study 1: longitudinal preterm and full-term infant fecal samples

#### Study overview

The first case study consists of 402 fecal samples collected from 19 preterm and 23 full-term infants from birth to age one, and their mothers around time of birth [11]. This study was designed to investigate strain persistence within infants, strain sharing between infants and their mothers, and strain sharing among different infants.

DNA extraction was primarily achieved using 96-well plates (Methods) and there were five extraction plates, labeled P1 to P5. There were five reagent-only negative controls, one for each DNA extraction plate, and they were labeled by the plate number (i.e., NC1 refers to the negative control on P1). One ZymoBIOMICS Microbial Community Standard (catalog #D6300; termed “Zymo”) was included as a DNA extraction positive control on P5. Following DNA extraction, all samples including controls were subjected to metagenomics sequencing and *de novo* genome reconstruction. Dereplication of the genomes constructed from fecal samples resulted in 1,005 representative genomes, as previously described [11]. Reads from all samples were then mapped to this dereplicated genome set for organism detection (Methods). To detect potential sources of contamination, we examined strain sharing among unrelated samples within and across extraction plates (Methods). Samples are considered unrelated if they are from different infants that are not biologically related or if one sample is a negative control.

#### Evaluation of extraction negative and positive controls

No organism was detected in negative controls NC1 and NC5. However, negative controls NC2, NC3, NC4 had at least one read mapped to ≥50% of at least one of the 1,005 representative genomes (this value served as our threshold for detection; Figure 1). No contaminants were detected in the Zymo positive control.

**Figure 1.**
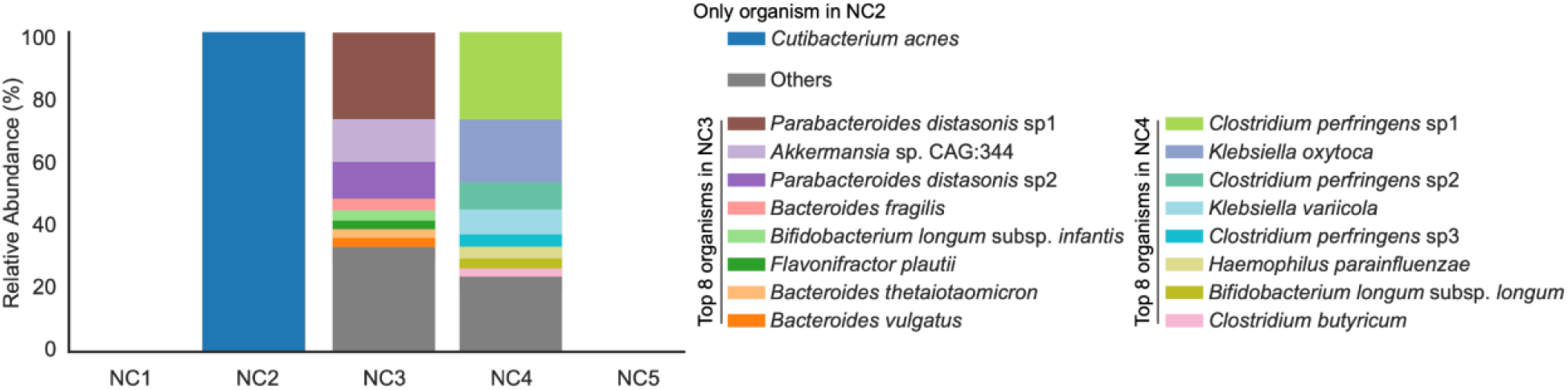
Microbial organisms were detected from three out of five negative controls. Eight most abundant genomes from each NC are colored in stacked bar plots. The rest of organisms are all grouped into “Others”.

The only genome detected in NC2 was *Cutibacterium acnes*. This organism was also found in nine fecal samples from five infants. These samples were extracted on P1, P2, and P4. Strain-to-strain comparisons indicate that all strains are sample-specific and different to that in NC2. Therefore, the presence of *C. acnes* in NC2 was not considered to be due to cross-sample contamination. As *C. acnes* is a common skin commensal bacterium [16], we suspect this organism is an externally derived contaminant.

NC3 and NC4 each had ~60 genomes that were above our read-mapping detection threshold. The organisms represented by these genomes could be externally derived and/or originated within the study via index switching, sample bleeding and/or well-to-well DNA contamination. If any of these genomes were derived from external sources (e.g., from DNA extraction kits), we would expect the strains to be in the majority of samples from the same plate (because they were processed simultaneously) and perhaps across extraction plates. However, no strain was shared among negative controls. Further, the strains found in NC3 and NC4 were only shared by a maximum of 7.7% and 6.5% of the unrelated samples from PC3 and PC4, respectively. Thus, we conclude that the genomes in NC3 and NC4 were unlikely due to external contamination and likely originated from other samples from this study.

#### Evaluation of cross-sample contamination: index-hoping and sample bleeding

The majority of strains found in NC3 and NC4 were only shared with samples from the same extraction plate. This observation rules out index switching and sample bleeding, as these phenomena should lead to contamination of samples from other plates because DNA from different plates was pooled for sequencing (P1 and P2 were pooled and sequenced at a different time than samples from P3-P5). Both index switching and sample bleeding refer to the misassignment of reads to samples. However, index switching results from indices being similar in multiplexing sequencing [8], and can be essentially prevented by using Unique Dual Indexes, which was what we used for sequencing in this study. Sample bleeding, on the other hand, occurs due to the close proximity of sample read clusters on the flow cell during sequencing [9]. We confirmed that this is not the main explanation for the contamination in NC3 and NC4 by resequencing these two controls and finding that the community compositions were essentially the same as the originally sequenced NC3 and NC4.

#### Evaluation of cross-sample contamination: well-to-well contamination

To test for the remaining possible explanation for the contamination in NC3 and NC4, well-to-well contamination, we visualized strain sharing patterns in the context of the extraction plates. Based on the observation made by Minich et al. that neighboring wells are more prone to well-to-well contamination than distant wells [7], we hypothesized that, if between-sample contamination occurred within an extraction plate, nearby unrelated sample pairs would more likely to share strains than those that were farther apart. Distance-correlated strain sharing was not seen on P1, P2 and P5 (p = 0.18, 0.31, and 0.32, respectively; Wilcoxon rank-sum test). This finding is consistent with the plate-based strain sharing visualization, which shows that unrelated samples on P1, P2 and P5 rarely shared any strains and for pairs that did share strains, the majority of them were not immediately adjacent to each other (**Figure 2**).

**Figure 2.**
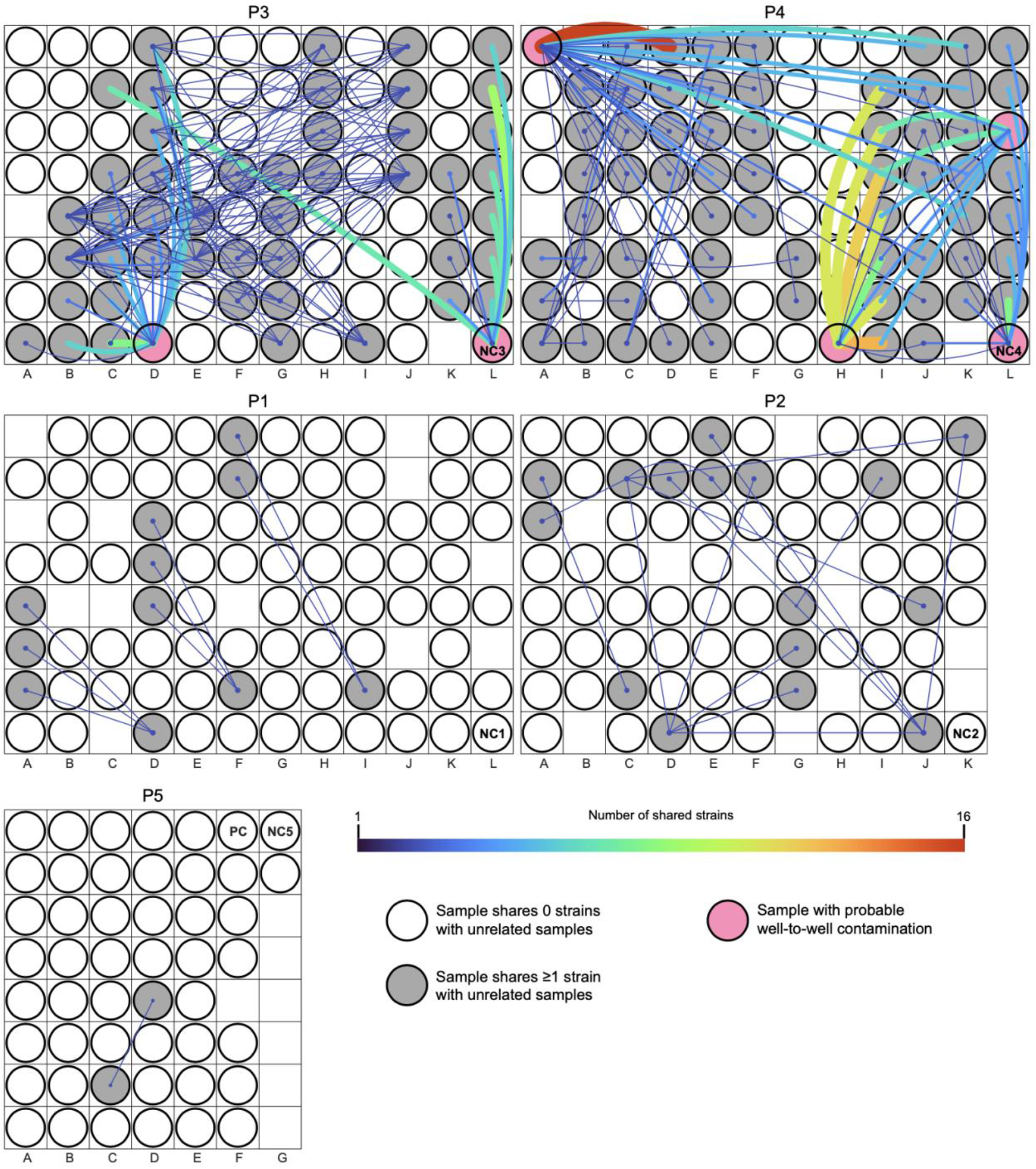
Within-plate strain sharing among unrelated samples. Rectangular areas represent plates (P1-P5) and circles show sample placements within each plate. A line was drawn between unrelated samples if they shared ≥1 strain. The more strains a sample pair shared, the thicker and brighter the line. If a sample did not share any strains with other unrelated samples, its corresponding circle is colorless. Pink circles represent samples that were likely cross-contaminated.

On P3 and P4, however, nearby samples were significantly more likely to share strains than those that were farther apart (p = 2.3e-3 and 4.7e-3, respectively; Wilcoxon rank-sum test). Plate-based strain sharing visualization indicates that a few samples including NC3 and NC4 in particular (pink circles in **Figure 2**; see also **Figure S1**) exhibited location-specific sharing patterns, consistent with well-to-well contamination. For instance, on P3, NC3 located on column L primarily shared strains with samples from infants #82 and #83 that were loaded onto columns K and L for extraction. This suggests that the fecal samples from infants #82 and #83 were potential sources of the contamination seen in NC3. NC3 also shared strains with a sample from infant #83 that was loaded onto column C for extraction. As expected, samples from infant #83 share strains, so strains in NC3 could have come from any of the samples from infant #83. Likely the contaminant strains were derived from the samples adjacent to the negative control. Thus, we do not attribute this instance of sharing to long distance well-to-well contamination. NC4 on P4 exhibited a similar proximity-based strain sharing pattern to NC3 (**Figures 2** and **S1**), and we deduce that NC4 was primarily cross-contaminated by adjacent samples.

In addition to NC3 and NC4, four preterm infant samples displayed plate location-specific strain sharing patterns (unlabeled four pink circles in **Figure 2**). The first instance involved a sample from infant #98 (termed #98D4). #98D4 was extracted on P4 and it shared strains with nearly half of the samples extracted from the same plate, including those on columns J and K that were nine and ten columns away (**Figure 2**). Shared strains were from six infants, including a pair of twins (#122 and #123). Strains shared by each of the twins and #98D4 overlapped completely and these strains were not found in any other infants. The other four of these six infants also shared strains that were otherwise unique to them with #98D4. #98D4 did not share strains with samples extracted from different plates. We therefore deduce that #98D4 was contaminated by multiple wells on P4. Besides #98D4, three other samples, each derived from a different infant (#63, #114 and #128), primarily shared strains with neighboring samples, similar to the patterns seen in NC3 and NC4. We confirmed that these four infant samples were indeed cross-contaminated by re-extracting and sequencing three of the four samples (the original stool sample from #128 and its close-by-date replacement were unavailable). For the sample from infant #63, a close-by-date sample (day of life 6 instead of 9) was selected as the original stool sample was unavailable. For all three re-extracted samples, their DNA concentrations became ~0 (**Table S1**) and their location-based strain sharing patterns were not observed.

#### Evaluation of underlying biological signals after removing contaminants

The identification of well-to-well contaminated samples allowed us to assess strain sharing among supposedly uncontaminated infant samples on originally discarded P3 and P4. On P3, samples from six preterm infants shared a total of four strains. These four are strains of *Clostridioides difficile, Clostridium paraputrificum, Clostridium butyricum*, and *Lactobacillus rhamnosus* (**Figure 3**). Samples that shared these four strains were not often adjacent to one another. Further, for each of these four organisms, near-identical strains were also found among samples that were extracted on different plates. Notably, *C. difficile, C. paraputrificum*, and *C. butyricum* strains were shared among preterm infants only. The pattern of strain sharing on P4 was similar to that on P3, except that there were more shared strains that were shared by fewer samples (11 strains were shared among samples from four infant pairs). As for P3, we detected no strong signal for contamination via dispersal of strains into all or most surrounding wells from single source wells. Given minimal evidence of well-to-well contamination and since all infants in this study were born in the same hospital, we conclude that most of the strains shared by the infants whose samples were extracted on P3 and P4 were probably hospital derived, a phenomenon that has been reported previously [17].

**Figure 3.**
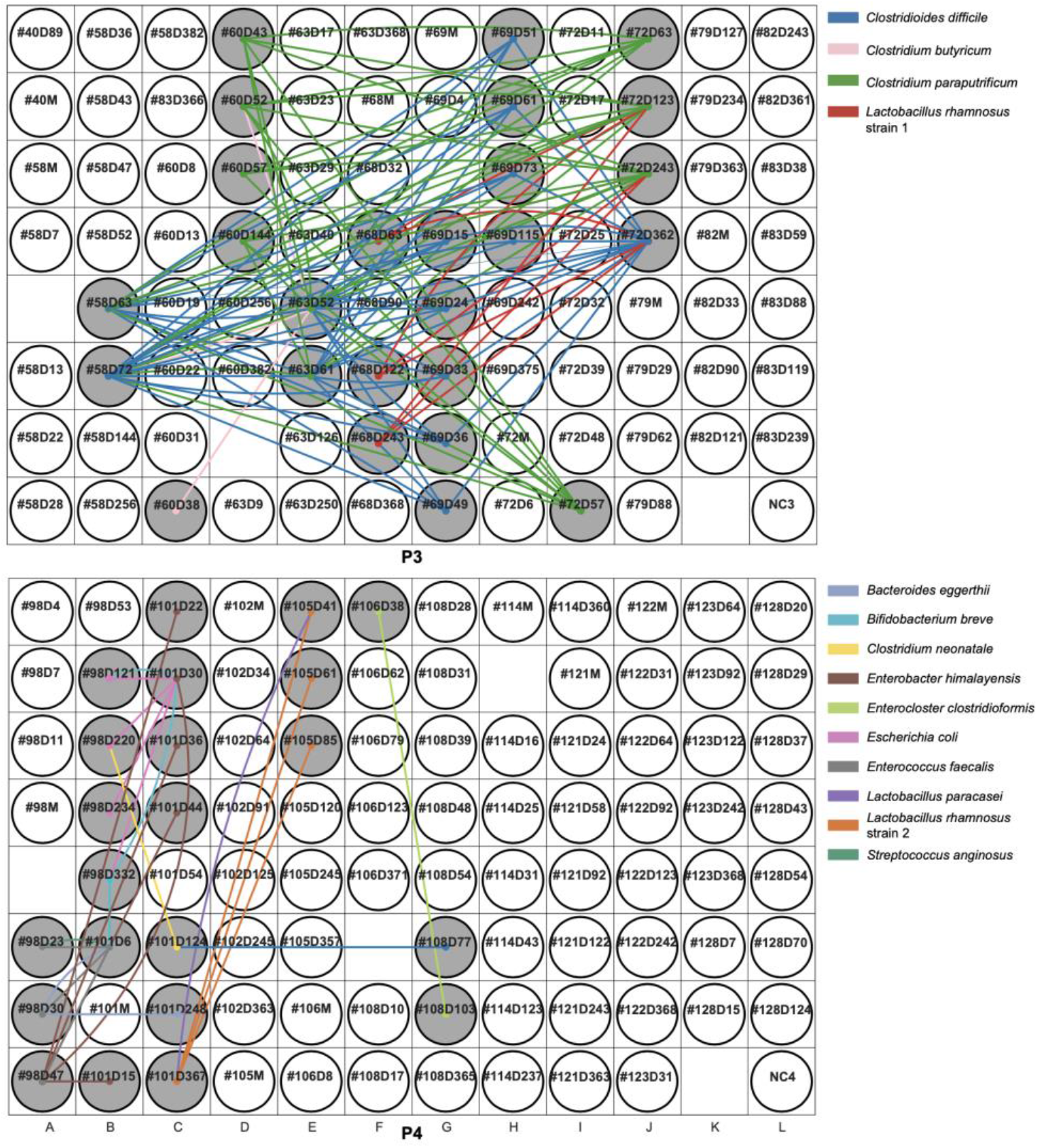
Strain sharing within P3 and P4 after removing cross-contaminated samples. Rectangular areas represent plates (P3, P4) and circles show sample placements within each plate. Infant samples are named by infant number and infant day of life (i.e., #40D89 refers to infant #40 and this sample was collected when the infant was 89-day-old). Maternal samples are named by the infant number with a letter “M” in the end (i.e., #40M refers to the maternal fecal sample collected from infant #40). A line was drawn between unrelated samples if they shared a strain and the line was colored by strain type. If a sample did not share any strains with other unrelated samples, its corresponding circle is colorless.

#### Conclusions from Case Study 1

Using strain-resolved workflow, we identified well-to-well contamination to be the major source of contamination in this dataset. The six contaminated samples (two negative controls and four preterm infant samples) were all low in microbial biomass. If such contamination was not addressed, we would have falsely concluded that strain sharing among non-related infants was not rare and that some non-related infant pairs could share as many strains as sibling pairs do.

### Case study 2: longitudinal preterm infant fecal, mouth and skin samples

#### Study overview

We applied our strain-resolved workflow to a different clinical dataset consisting of 533 samples collected from the skin, mouth and stool of 16 preterm infants. These preterm infants were born in the same hospital as the infants from case study I. This dataset was part of a study designed to elucidate strain transmissions between the hospital environment and preterm infants.

DNA extraction was primarily achieved using 96-well plates. One reagent-only negative control was included in each extraction plate. There were six plates (labeled P1 to P6) and six negative controls, labeled NC1 to NC6. In addition, P4, P5 and P6, each had ~3 Zymo standards as DNA extraction positive controls.

236 of the 533 samples (including 3 negative controls (NC1, NC3 and NC5) and 3 positive controls, one from P5 and two from P6; termed PC5, PC6_1, and PC6_2) were selected for metagenomics sequencing (colored circles except for those light blue ones in plate maps in **Figures 5-7**). Before library preparation, DNA was transferred from the extraction plates to three new 96-well plates, one for each sample type (Methods). Following sequencing, d*e novo* genome reconstruction and dereplication yielded 152 representative genomes, which served as the reference genomes for read-mapping based organism detection for this dataset (Methods). To detect potential sources of contamination, we examined strain sharing among all unrelated samples, as described in case study I.

#### Evaluation of extraction negative and positive controls

Of the three sequenced negative controls, one genome was detected in NC1 and NC3, and 9 genomes were detected in NC5 (**Figure 4**). No contaminants were detected in the Zymo positive controls.

**Figure 4.**
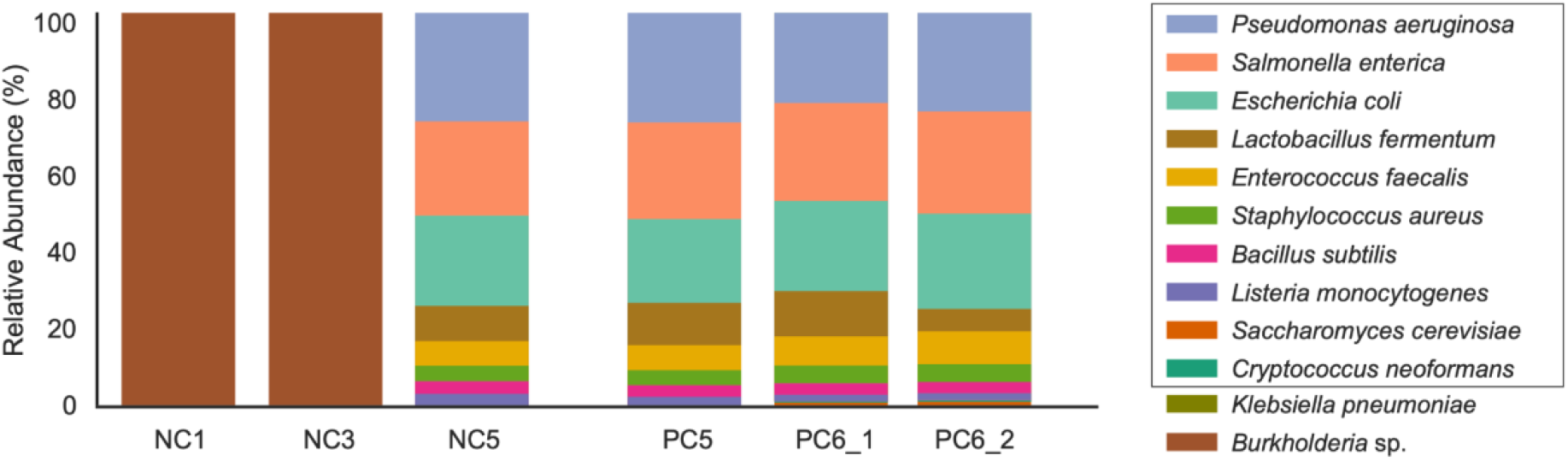
Community composition of negative and positive controls. Relative abundance of genomes in the negative and positive controls (NC and PC, respectively), colored by organism type. The first 10 organisms listed (boxed) are those in the Zymo standard.

*Burkholderia* sp. was the only organism detected in NC1 and NC3. The *Burkholderia* strain in NC1 was not detected in any sequenced samples. However, a different *Burkholderia* strain was found in NC3 and 12 infant skin samples that were extracted from four extraction plates (**Figure 5**). No *Burkholderia* was found in fecal or mouth samples, both of which were higher in biomass than skin samples (p = 5.38e-37 and 1.58e-40, respectively; Wilcoxon rank-sum test). The skin samples that contained the *Burkholderia* strain had a significantly lower biomass than the skin samples that did not (p = 2.79e-24; Wilcoxon rank-sum test). *Burkholderia* is not part of the normal skin microbiome [16], but it has often been reported as a reagent contaminant [1,18–20]. Identifying one single *Burkholderia* strain among infant and negative samples suggests this strain is a result of external contamination. Interestingly, this strain is also in one low-biomass gut sample from case study I. This contaminant was likely introduced prior to library preparation as it was not detected in NC1 and NC5, both of which were on the same library preparation plate as those *Burkholderia*-containing skin samples **(Figure 5C)**.

**Figure 5.**
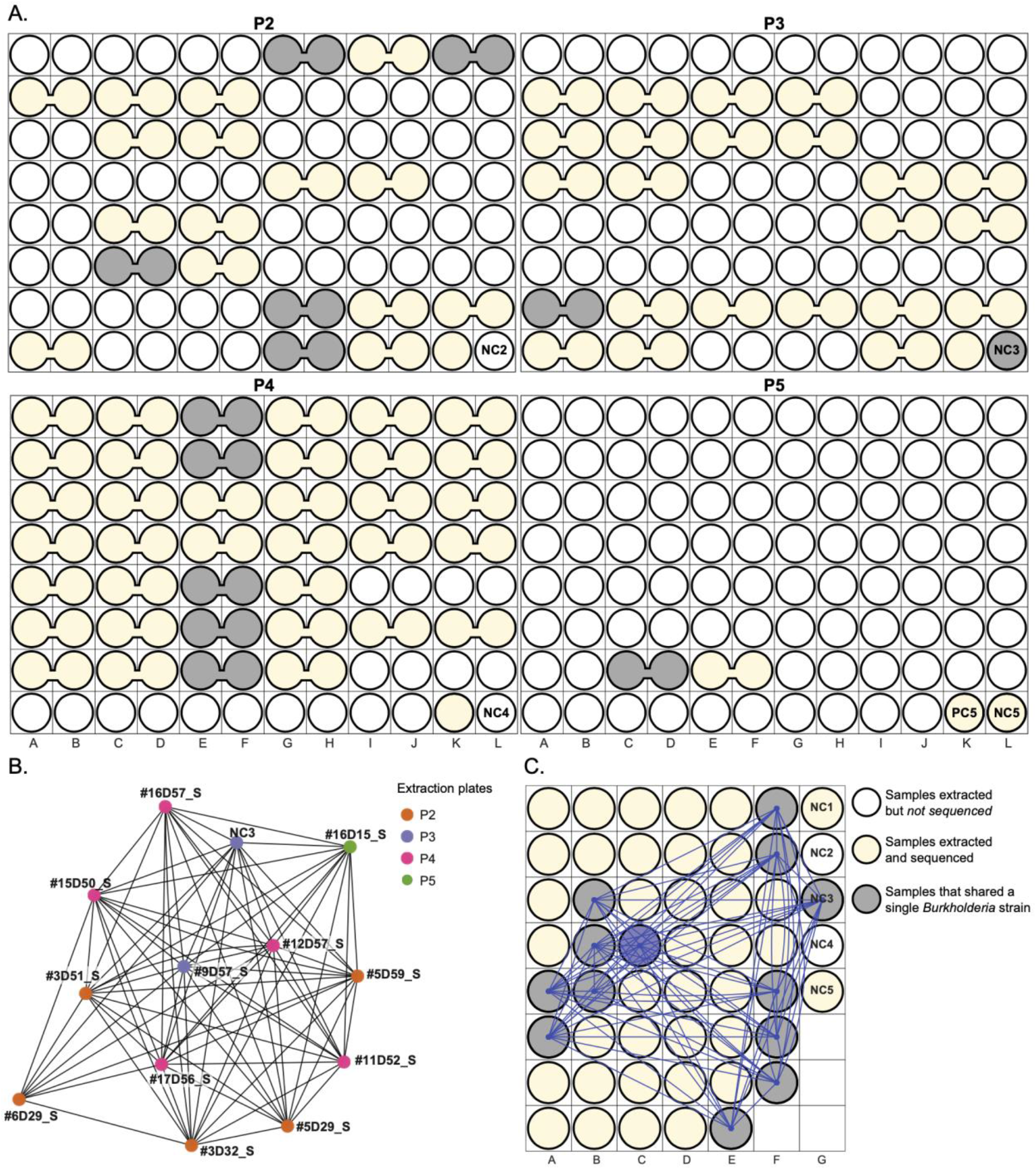
A single *Burkholderia* strain is shared across 13 samples. Samples (circles) that were sequenced are yellow and those that contain a *Burkholderia* strain are gray. The remaining colorless circles represent samples that were extracted but not sequenced. A) Extraction plate locations of 13 samples that shared a single *Burkholderia* strain. Merged circles represent duplicated samples that were extracted adjacent to each other and were merged before being transferred to the library preparation plates. B) *Burkholderia* strain sharing network. Each node represents a sample and is colored by the extraction plate. Nodes are connected if they share the *Burkholderia* strain. Infant samples are named by infant number, infant day of life and sample type (“M” refers to mouth samples, “S” refers to skin samples, and “G” refers to gut samples). For instance, #16D57_S refers to infant #16’s skin sample and this sample was collected when the infant was 57 days old. C) Library preparation plate displaying the location of samples that shared a *Burkholderia* strain. Lines were drawn between circles if their corresponding samples shared the *Burkholderia* strain.

For NC5, one organism, *Klebsiella pneumoniae*, was present at an extremely low abundance (<0.1%) and was not detected in any other samples. We cannot determine if adjacent samples on the extraction plate were possible sources of this organism because those samples were not sequenced. The remaining 8 organisms in NC5 were all bacterial members of the Zymo community (**Figure 4**). NC5 was adjacent to PC5, a Zymo positive control, on the extraction plate. Thus, Zymo organisms in NC5 could be attributed to well-to-well contamination.

#### Evaluation of Zymo contamination in infant samples

To further evaluate contamination by the Zymo strains, we searched for these strains in infant-derived samples. During DNA extraction, six skin and oral samples were deliberately spiked with 75 μL of the Zymo standard, four of which were sequenced (Methods). By examining strains shared between positive controls and biological samples, we found 12 additional infant samples containing at least the four most abundant Zymo members (**Figure 6**). All but one of these samples were from skin or mouth, which had lower biomass than gut samples (p = 1.58e-40 and 5.57e-9, respectively; Wilcoxon rank-sum test). Of these 12 infant samples, 9 were adjacent to a Zymo spiked infant sample or a positive control. Since the contaminated samples generally only shared Zymo strains with neighboring Zymo spiked samples (and not the other organisms in those samples), we conclude that the observed Zymo contaminants were more likely to be introduced accidentally, possibly via aerosolization or mis-pipetting, rather than via well-to-well contamination.

**Figure 6.**
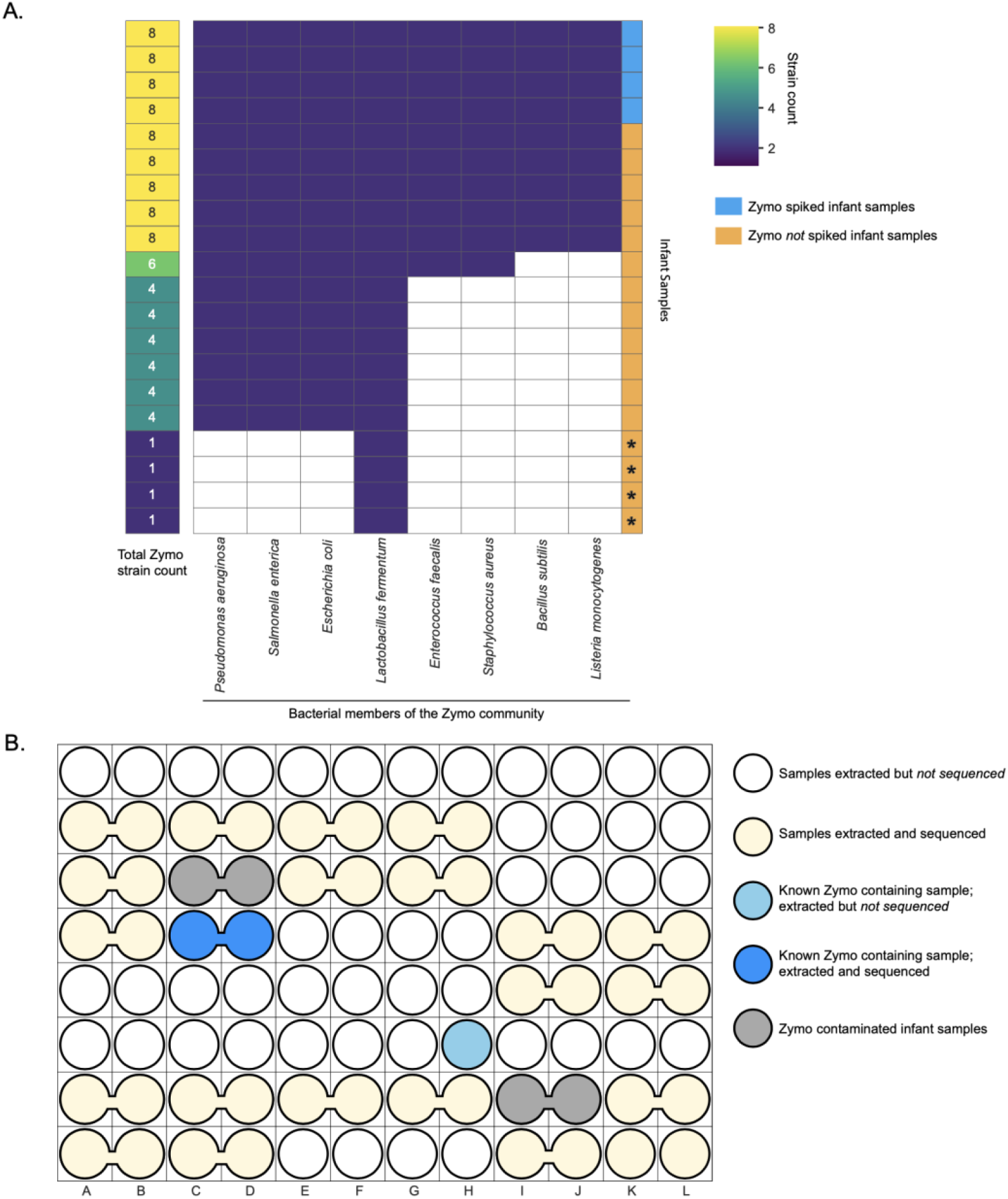
12 infant samples were contaminated with Zymo DNA. A) Clustergram displays the sharing of Zymo-associated strains between infant samples and Zymo positive controls. Each row corresponds to an infant sample that had ≥1 Zymo strain. The first column on the left displays the number of Zymo strains in each infant sample. The remaining 8 columns correspond to the 8 Zymo bacteria (from left to right, these 8 bacteria were arranged from the most to the least abundant) and they were colored by presence (dark purple) and absence (colorless) of the corresponding bacterium within each infant sample. “*” at the bottom four rows refers to samples that are not contaminated by the Zymo standard. B) A representative extraction plate (P3) showing two Zymo contaminated samples (gray bubbles). Merged circles represent duplicated samples that were merged post DNA extraction.

#### Evaluation of additional contamination not present in negative controls

Using the strain-resolved approach developed in case study I, we evaluated strain sharing among infants after excluding Zymo and *Burkholderia* strains. Five bacterial strains were widely shared by samples from different infants. Specifically, for each of these five strains, at least half of the infants had one sample that shared such a strain with a sample from an unrelated infant. Of these five strains, two are *Staphylococcus epidermidis* strains A and B, and the other three are *Staphylococcus* M0480, *Corynebacterium aurimucosum*, and *Cutibacterium acnes*. All five are common members of the healthy skin microbiome [16]. *S. epidermidis* strain A and the *Staphylococcus* M0480 strain were shared among all sample types (skin, mouth and stool), and *S. epidermidis* strain B, the *C. aurimucosum* strain and the *C. acnes* strain were shared among skin samples only. Additionally, a near-identical *S. epidermidis* strain A was found in fecal samples from 16 out of 42 infants from case study I. It is uncommon to find a single strain of each of these organisms in the majority of infants of a single dataset, we therefore identify these five strains to be externally-derived contaminants (e.g., from staff who handled the samples).

We re-examined strain sharing among samples of this dataset after excluding all identified external contaminants (Zymo strains, a *Burkholderia* strain, and five skin-associated strains). While most of the extraction plates did not exhibit location-based strain sharing, samples from one infant pair on P4 did, suggesting that there might be well-to-well contamination (pink circle in **Figure S2**). A sample from infant #12 shared up to 3 strains with neighboring skin and oral samples of infant #13. These 3 shared strains were not shared by infant #13 and any other infants. In addition, none of the other samples from infant #12 shared strains with infant #13. This suspected cross-contaminated infant #12 sample was collected from skin and its strain sharing pattern was similar to those of well-to-well contaminated samples in case study I.

#### Conclusions from Case Study 2

Our strain-resolved workflow identified external contamination to be the major source of contamination in this dataset. Suspected contaminants include *Burkholderia* strains, Zymo DNA, and five skin-associated strains. Our approach also suggested one skin sample to be cross-contaminated by adjacent samples from the same extraction plate. Notably, most of these contaminants were found in skin samples, which had lower biomass than oral and gut samples.

## DISCUSSION

Contamination is an insidious and potentially unavoidable problem in metagenomics-based microbiome research. If not appropriately addressed, extraneous DNA sequences can skew conclusions, resulting in potentially false statements. Here, we devised a workflow that relies on strain-resolution to detect contamination based on unexpected sharing of essentially identical strains, and demonstrate its usefulness in two clinical metagenomics datasets. By examining strain sharing based on genotype distribution across samples, and considering sample proximity on DNA extraction plates, we identified contamination that derived from external sources and that which originated within the sample set.

In a recent review, it was noted that negative controls have been included in ~30% of prior microbiome studies, although only a subset of studies sequenced and analyzed data from the negative controls [21]. Although negative controls can identify foreign DNA that does not belong to the study, they only offer a limited view of contamination and do not constrain the contamination source or the number of samples impacted. In our first case study, analysis of the pattern of shared strains in the context of sample location on DNA extraction plates indicated that the DNA in negative controls was mostly derived from neighboring biological samples. Detection of contamination motivated a more complete analysis of well-to-well contamination in all biological samples in our study. In four cases, genotypes found in many samples from one infant were shared by only one of the samples from another infant. In each such case, the contaminated sample was located adjacent to the putative contamination source on the DNA extraction plate. This conclusion was verified, as the strains were no longer shared when the DNA was re-extracted and resequenced. In our second case study, two externally derived contaminants in the negative controls were identified: *Burkholderia* strains and DNA from the Zymo positive control. We thus investigated strains that were apparently shared by samples from unrelated infants, and identified DNA from five additional contaminants in samples from the majority of infants yet absent in the negative controls.

It may be possible to find true biological signals if the signal from contamination can be removed. In case study one, after removing the well-to-well contaminated samples, we identified three Clostridia strains that were shared only among preterm infant samples. Since preterm infants in our study spent their first 2-3 months in the hospital, they have a higher chance to pick up hospital-associated strains than full-term infants. Given that strains of these bacteria have been found previously in preterm infants that were born in the same hospital and in neonatal intensive care unit room microbiomes [17,22], we hypothesize that these three strains may be circulating among infants in the hospital.

By identifying contaminants and estimating their origins, our study provides a detailed workflow for contamination identification. Based on our observations and previous contamination-related studies, we list several suggestions for minimizing contamination in metagenomics-based studies. First, we encourage others to minimize plate-based extraction if possible. If plate-based extraction is a must, one should take additional precautions such as limiting the number of open wells when extracting, using individual caps for covering wells on the plate and fully spinning down the samples before removing the caps to prevent well-to-well contamination from occurring. In addition, one should include their DNA extraction plate maps in their published work. Second, we urge others to randomize samples when extracting DNA. Specifically, one should avoid loading samples from the same individual or experimental group or biologically related individuals adjacent to one another for extraction because if well-to-well contamination occurs, such arrangements can blur the line between genuine signals and contamination and artificially inflate metagenome similarities among samples that were nearby. In our original publication using the dataset presented in case study I [11], when assessing persisting infant gut strains, we decided to not use any samples extracted from the two extraction plates in which cross-contamination occurred because we could not determine the degree of well-to-well contamination among samples from the same individuals. Third, sequencing of sampling negative controls (i.e., empty swab that is used during sample collection) is recommended in addition to extraction negative controls. This should identify contamination introduced during sampling, which is important because such contaminants will likely not display extraction-plate-based sharing patterns. Removal of strains seemingly shared but introduced during sampling will clarify strains truly shared by infants.

### Conclusion

Contamination may be unavoidable in high-throughput sequencing and our results suggest that it can be particularly problematic for low-biomass samples. Genotype-level surveillance has the advantage that it does not require additional expenditures related to library construction and sequencing. Our work emphasizes the importance of routinely assessing contamination prior to data analysis so as to avoid incorrect findings. As microbiome-based analysis and diagnosis are becoming more popular, we conclude that use of genotype-specific surveillance methods as well as negative controls to ensure the integrity and reproducibility of the results.

## Additional Files

### Additional File 1: Figure S1. Details of strain sharing on P3 and P4 from case study I

Rectangular areas represent plates (P3 and P4) and circles show sample placements within each plate. Infant samples are named by infant number and infant day of life (i.e., #63D9 refers to infant #63 and this sample was collected when the infant was 9-day-old). If it is a maternal sample, such a sample is named by the infant number with a letter “M” in the end (i.e., #40M refers to the maternal fecal sample collected from infant #40). A line was drawn between unrelated samples if they shared ≥1 strain. The more strains a sample pair shared, the thicker and brighter the line. If a sample did not share any strains with other unrelated samples, its corresponding circle is colorless. Pink circles represent samples that were likely cross-contaminated.

### Additional File 2: Figure S2. Detection of one cross-contaminated sample on P4 from case study II

Merged circles represent duplicated samples that were extracted adjacent to each other and were merged before being transferred to the library preparation plates. Infant samples are named by infant number, infant day of life and sample type (“M” refers to mouth samples, “S” refers to skin samples, and “G” refers to gut samples). If a sample pair from unrelated infants shared ≥1 strain, the corresponding samples circles were colored grey and a line was drawn between them. The more strains a sample pair shared, the thicker and brighter the line.

### Additional File 3: Table S1. The original and the re-extracted DNA concentrations of four cross-contaminated samples from case study I

*: The re-extracted stool sample (#63D6) is not the same as the original one (#63D9) as the original stool sample is unavailable. This alternative stool sample was collected 3 days earlier than the original sample.

## Declarations

### Ethics approval and consent to participate

This work was reviewed and approved by the University of Pittsburgh Institutional Review (IRB STUDY19120040 and PRO16050140).

### Consent for publication

All authors in this study have reviewed the results, read the manuscript, and gave their consent for publication.

### Availability of data and materials

The datasets supporting the conclusions of this article are included within the article and its additional files.

### Competing interests

JFB is a cofounder of Metagenomi. The remaining authors declare that they have no competing interests.

### Funding

This research was supported by the National Institutes of Health (NIH) under award RAI092531A and the Alfred P. Sloan Foundation grant 20110-6-06. This work used QB3 Vincent J. Coates Genomics Sequencing Laboratory at UC Berkeley, supported by NIH S10 OD018174 Instrumentation Grant.

### Authors’ contributions

YCL and JFB designed the study; MJM and BAF coordinated the sample collection; BAF performed sample DNA extractions; YCL and MRO coordinated the acquisition of the metagenomics data; YCL and JH performed the data analysis; JWR and SD assisted in the analysis and consulted on statistical methodology; YCL and JFB wrote the manuscript. All co-authors read and approved the manuscript.

## Acknowledgements

We thank Rohan Sachdeva, Ka Ki Lily Law and Shufei Lei for the technical support. For funding support, we acknowledge NIH award RAI092531A and Sloan grant to JFB and MJM and Chan Zuckerberg Biohub support to JFB.

## Methods

### Sample collection

Infant fecal samples from case study I were collected either at UPMC Magee-Womens Hospital by trained nurses or at home by parents provided with detailed collection instructions. Sample collection and storage details see Lou et al. [11].

For case study II, all infant samples (skin, oral and stool samples) were collected at UPMC Magee-Womens Hospital by trained nurses. Skin and oral samples were obtained by the charge nurse using a BD BBL Culture Swab EZ under supervision of study personnel. Skin and oral samples were collected in duplicate at each timepoint for each preterm infant in order to increase the biomass available for DNA extraction. Details of sample collection and storage see Olm et al. [23].

### DNA extraction

DNA was extracted using either the Qiagen QIAamp PowerFecal Pro DNA Isolation kit (single tube extractions; used for 14 of 402 samples in case study I and 7 of 533 samples in case study II) or Qiagen DNeasy PowerSoil HTP 96 DNA Isolation kit (96-well plate extractions; used in the majority of samples in both case studies) with modifications to the manufacturer’s protocol. To minimize cross-plate contamination, no plates were extracted at the same time. For each 96-well DNA extraction plate, a reagent-only negative control was included. ZymoBIOMICS Microbial Community Standard (catalog # D6300) was included as a positive control on one extraction plate from case study I and three extraction plates from case study II. When loading samples into the wells of DNA extraction plates, samples were not randomly distributed among the plates, and often, samples from the same infant were present next to each other sequentially along columns on the same extraction plate.

For DNA extracted from stool using the single tube format, the manufacturer’s protocol was followed except for a heating step at 65°C for 10 minutes before the homogenization step. For DNA extracted from stool with the 96-well kit, fecal samples were added to individual wells of the bead plate and stored overnight at −80°C. The following day, the Bead Solution and Solution C1 were added, and the plates were incubated at 65°C for 10 minutes. The plates were shaken on a Retsch Oscillating Mill MM400 with 96-well plate adaptors for 10 minutes at speed 20. The plates were rotated 180° and shaken again for 10 minutes at speed 20. All remaining steps followed the manufacturer’s centrifugation protocol.

All skin and oral samples from case study II were extracted using 96-well plates. Specifically, for skin and oral swab samples, the swab head was cut off directly into the wells of the bead plate and stored overnight at −80°C. The following day, the Bead Solution and Solution C1 were added, and the plates were incubated at 65°C for 10 minutes. The plates were shaken on a Retsch Oscillating Mill MM400 with 96 well plate adaptors for 5 minutes at speed 20. The plates were rotated 180° and shaken again for 5 minutes at speed 20. The Solution C2 and C3 steps were combined (200 μl of each added) to improve DNA yield. All remaining steps followed the manufacturer’s centrifugation protocol. For six selected skin and oral samples, 75 μl of ZymoBIOMICS Microbial Community Standard (catalog # D6300) was added to the wells of these six samples prior to the heating step during DNA extraction.

Extracted DNA was quantitated using the Quant-iT High Sensitivity dsDNA Assay Kit (Thermo Fisher Scientific) in 96-well plates and measurements made on a SpectraMax M2 microplate reader. All extracted samples from case study I were sequenced whereas only about half of the extracted samples from case study II were sequenced.

DNA extractions and quantifications were performed at the University of Pittsburgh School of Medicine, Pittsburgh, PA. Once completed, the extracted DNA was shipped to the QB3 Vincent J. Coates Genomics Sequencing Laboratory at UC Berkeley for library preparation and sequencing. For case study I, the extracted DNA was sent in the same plates in which the DNA was eluted in the final step of the DNA extraction protocol. For case study II, the DNA from selected samples was transferred from the extraction plates to three new 96-well plates (one for skin samples, one for oral samples, and one for stool samples) before shipping to Berkeley. For each pair of the duplicated skin and oral samples, their extracted DNA was combined into a single volume on the new 96-well plates.

### Metagenomic sequencing

Samples from case study I and II had separate library preparation and sequencing runs. However, the overall sequencing workflow is essentially identical. Metagenomic sequencing of all samples was performed in collaboration with the California Institute for Quantitative Biosciences at UC Berkeley (QB3-Berkeley). Library preparation on all samples from each study was performed as previously described [24]. Final sequence ready libraries were visualized and quantified on the Advanced Analytical Fragment Analyzer. All libraries were then evenly pooled into a single pool and sequenced on individual Illumina NovaSeq6000 150 paired-end sequencing lanes with 2% PhiX v3 spike-in controls. Post-sequencing bcl files were converted to demultiplexed fastq files per the original sample count with Illumina’s bcl2fastq v2.20 software.

### Metagenomic assembly, *de novo* binning, and taxonomy assignment

Sequencing reads from case study I and II were assembled separately. However, the overall workflow of metagenomics assembly, de novo binning and taxonomy assignment was essentially identical. See Lou et al. for details on read assembly, *de novo* binning, and taxonomy assignment on resulting genome bins [11].

### Genome dereplication

To generate a set of study-specific, high-quality, and nonredundant reference genomes, all de novo assembled genome bins were dereplicated at 98% whole-genome average nucleotide identity (gANI) via dRep (v2.6.2) [25], using a minimum completeness of 75%, maximum contamination of 10%, the ANImf algorithm, 98% secondary clustering threshold, and 25% minimum coverage overlap. Since biological samples from case study II were deliberately spiked with Zymo standard (catalog #D6300), 10 publicly available Zymo genomes (https://s3.amazonaws.com/zymo-files/BioPool/ZymoBIOMICS.STD.refseq.v2.zip) were added to the original genome set of case study II before dereplication. Genomes with gANI ≥98% were classified as the same “subspecies”, and the genome with the highest score (as determined by dRep) was chosen as the representative genome from each subspecies.

### Detection of subspecies and identification of strains using inStrain

Reads from each individual sample were mapped to study-specific representative subspecies (generated via dRep as described above) concatenated together using Bowtie2 under default settings. inStrain (v1.3.4) *profile* [26] was run on all resulting mapping files using a minimum mapQ score of 0 and insert size of 160. Genomes with ≥0.5 breadth (meaning at least half of the nucleotides of the genome are covered by ≥1 read) in samples were considered to be present. inStrain *compare* was used under default settings to compare read mappings to the same genome in different pairs of samples. In case study I, samples were considered to share the same strain of the examined genome if the compared region of the genome from samples shared ≥99.999% population-level ANI (*popANI*) whereas in case study II, the popANI threshold was set to be 99.995%. Only genomic areas with at least 5x coverage in samples were compared, and sample pairs with less than 50% of comparable regions of the genome were often excluded (≥0.5 *percent_genome_compared*). For edge cases, such as when *popANI* values were within 0.005% of the threshold or when *percent_genome_compared* values were within 0.2% of the threshold, inStrain *compare* results were manually assessed to determine whether the sample pairs shared the same or different strains.

### Statistical analysis

Statistical significance for two-group univariate comparisons was calculated using Wilcoxon rank-sum test (as implemented using the Scipy module “scipy.stats.ranksums”) as reported in the main text. For instance, to assess whether strain sharing was correlated with sample pair distance on each DNA extraction plate, we compared within-plate Euclidean distances of sample pairs that did not share strains to those that shared at least one strain using Wilcoxon rank-sum test.

