## Supplementary figures and images for "Using strain-resolved analysis to identify contamination in metagenomics data"

### Figure S1

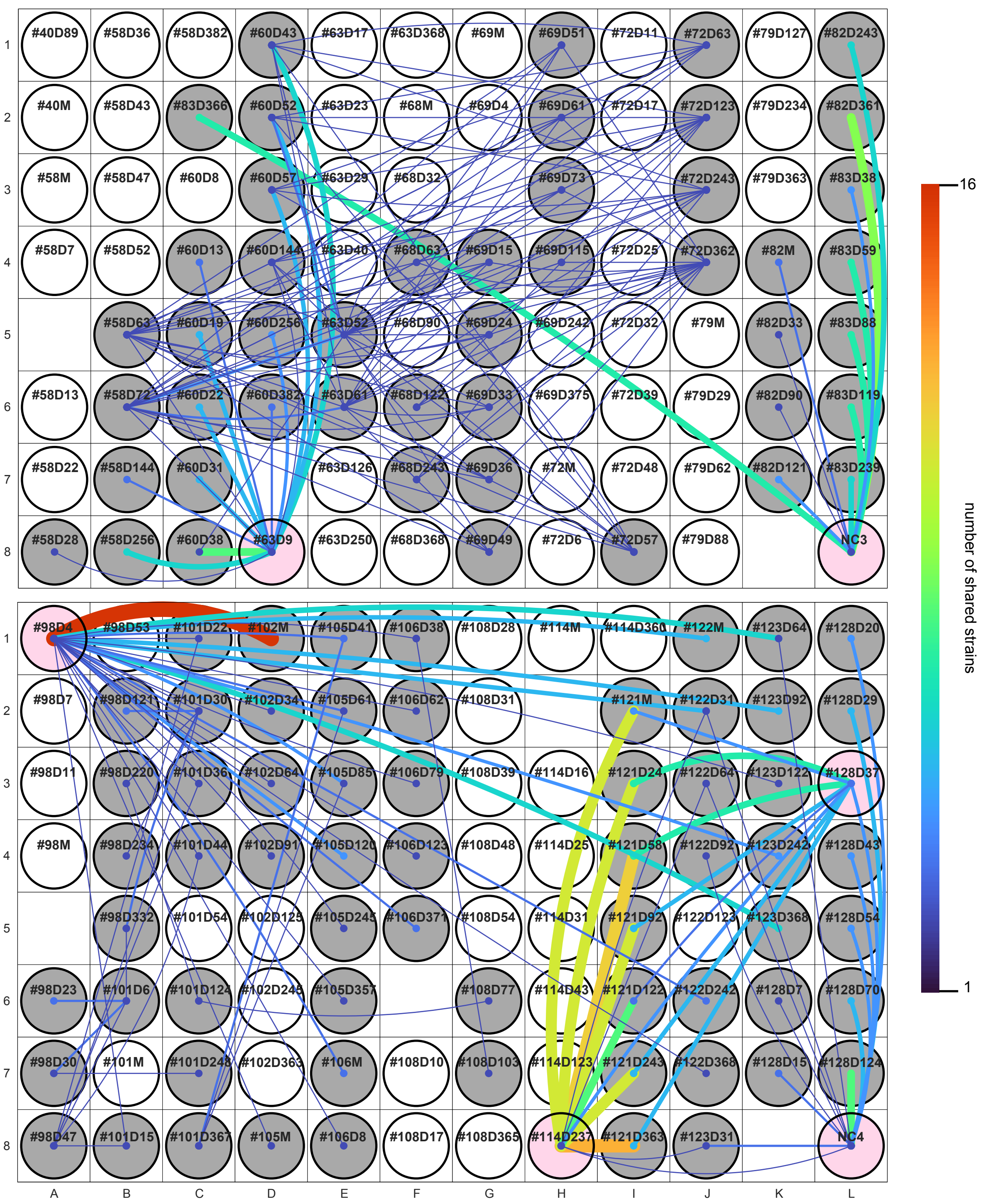

### Figure S2

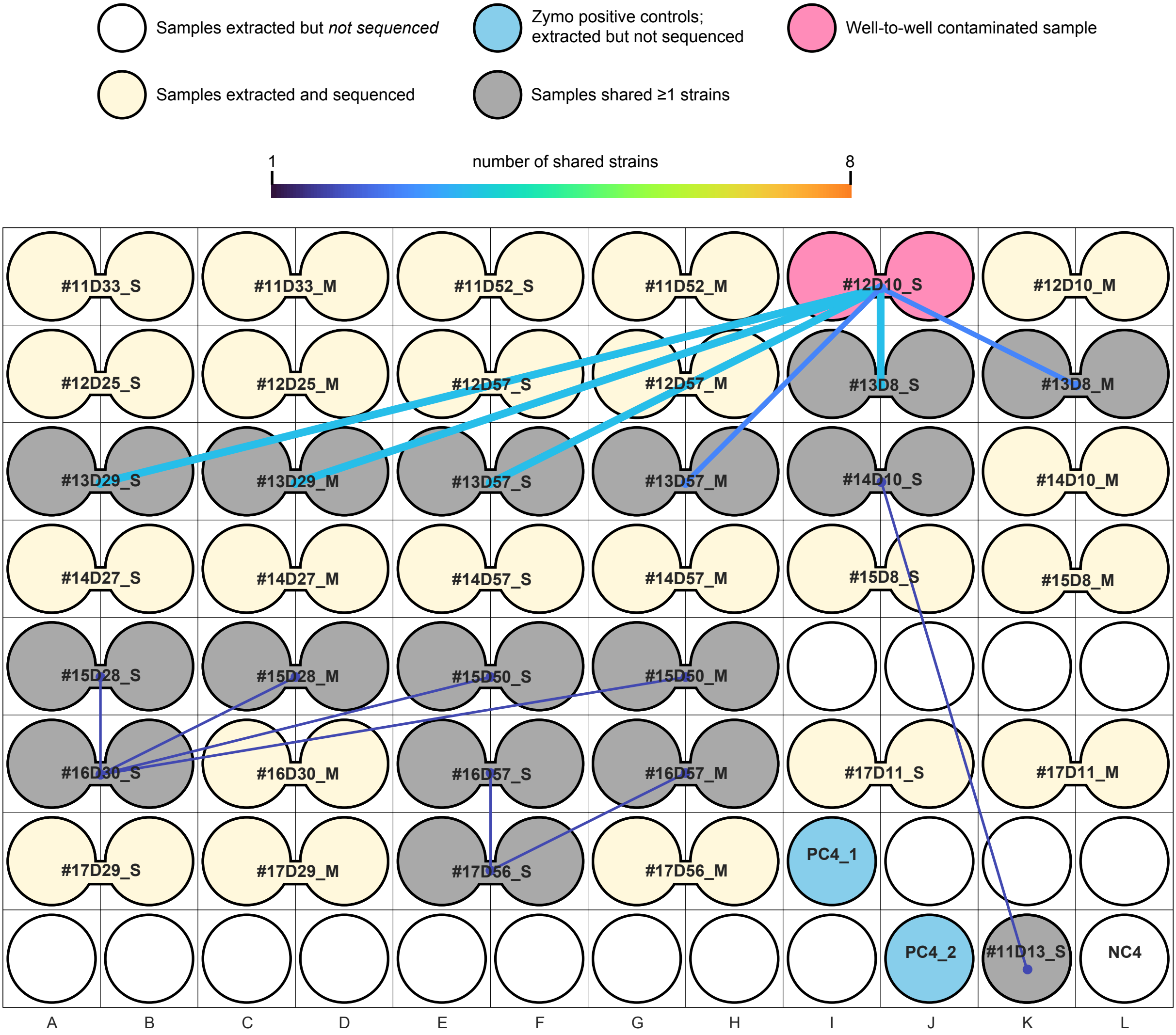
